# Unique Features of Sub-Cortical Circuits in A Macaque Model of Congenital Blindness

**DOI:** 10.1101/609636

**Authors:** Loïc Magrou, Pascal Barone, Nikola T. Markov, Gwylan Scheeren, Herbert P. Killackey, Pascale Giroud, Michel Berland, Kenneth Knoblauch, Colette Dehay, Henry Kennedy

## Abstract

There is extensive modification of the functional organization of the brain in the congenital blind human, although there is little understanding of the structural underpinnings of these changes. The visual system of macaque has been extensively characterized both anatomically and functionally. We have taken advantage of this to examine the influence of the congenital blindness in macaque resulting from the removal of the retina during in utero development. Developmental anophthalmia in macaque effectively removes the normal influence of the thalamus on cortical development leading to an induced hybrid cortex (HC) combining features of primary visual and extrastriate cortex. Here we show that retrograde tracers injected in early visual areas including hybrid cortex reveals a drastic reduction of cortical projections of the reduced lateral geniculate nucleus. In addition, there is an important expansion of projections from the pulvinar complex to the hybrid cortex, compared to the controls. These findings show that the functional consequences of congenital blindness need to be considered in terms of both modifications of the inter-areal cortical network and the ascending visual pathways.

## Introduction

There is a major functional reorganization of the cortex during development in the congenitally blind brain. Braille reading has been shown to activate the primary visual cortex, area V1, in early blind individuals (Sadato et al. 1996; Cohen et al. 1997; Buchel et al. 1998). In addition, the visual cortex of the congenitally blind has been observed to support higher cognitive functions including language and numerical processing (Cohen et al. 1997; Roder et al. 2002; Amedi et al. 2004; Bedny et al. 2011; Burton et al. 2012; Watkins et al. 2012; Kanjlia et al. 2016; Crollen et al. 2019). It has been hypothesized that such functional reorganizations reflect a global metamodal cortical function, where computations are defined by a particular local network, and where the reorganized brain function reflects a conserved cortical structure accommodating the changes in the inputs from the sensory periphery. Hence according to this theory, developmental deafferentation could induce a higher cognitive function in an early area of the visual cortex by the accommodation of possibly innately determined, long-distance inputs (Pascual-Leone and Hamilton 2001; Sur and Leamey 2001; Merabet and Pascual-Leone 2010; Bedny 2017; Amalric et al. 2018).

In contrast to the metamodal theory there is ample evidence form invasive animal experimentation that deafferentation during development leads to widespread changes in the adult brain. While the plastic changes that underpin cross-modal compensation are not fully understood it is generally agreed that they engage activity-dependent processes during brain development (Bavelier and Neville 2002; Ruthazer and Cline 2004). Further, there is strong evidence of a reorganization of the large-scale network in the cortex. Whole-brain networks of cortical thickness in sighted and early blind individuals suggests that the congenital blind exhibit an extensive reorganization of anatomical networks defined in this manner (Hasson et al. 2016). More recently this issue has been addressed in a macaque model of congenital blindness where anopthalmia is surgically induced in utero by bilateral removal of the retina (Magrou et al. 2018). This model of congenital blindness builds on earlier investigations which explored the impact of sensory input from the retina on the specification of the primary visual cortex, and where the absence of the retina during development led to part of the primary visual cortex acquiring profound changes in its cytoarchitecture to form a hybrid cortex (Rakic 1988; Dehay et al. 1989). Investigation of the cortical network in this model of congenital blindness showed a six-fold expansion of the spatial extent of local connectivity in the hybrid cortex and a surprisingly high location of the hybrid cortex in a computational model of the cortical hierarchy (Magrou et al. 2018). In the anophtalmic, the set of areas projecting to the hybrid cortex, areas V2 and V4 does not differ from that of normal adult controls, but there is a highly significant increase in the relative cumulative weight of the ventral stream areas input to the early visual areas. These findings show that although occupying the territory that would have become primary visual cortex, the hybrid cortex exhibits features of a higher order area, thus reflecting a combination of intrinsic and extrinsic factors on cortical specification (Rakic 1988; Dehay et al. 1989). Understanding the interaction of these contributing factors will shed light on the neurobiology of blindness.

Are these changes in the cortical network solely responsible for the functional reorganization observed in congenital blindness? Intrigued by the changes in dimensions of the thalamic nuclei in the anophtalmic monkey (Rakic et al. 1991; Dehay et al. 1996), in the present study we have examined the subcortical projections to the cortex of anophtalmic monkey. This revealed a massive reduction of the projections from the lateral geniculate nucleus to the hybrid cortex. This reduction of afferents from the major thalamic relay of the visual system was counterbalanced by a significant increase in projections from the pulvinar complex to the hybrid cortex. The marked changes from the pulvinar to the cortex contrasted with the observations that projections from other nuclei including the intralaminar thalamic nuclei, the amygdala and the claustrum were similar across experimental and control cases.

## Materials and Methods

We examined the connectivity of the cortex in two 25-day old and one 10-month-old macaque that had undergone retinal ablation at E58) and E73 (**Table 1**). In these three experimental anophtalmic animals, we made six tracer injections, a fast blue (FB) and a diamidino (DY) injection in each and we compared the results to 10 injections made in 8 adult controls.

**Table 1.**
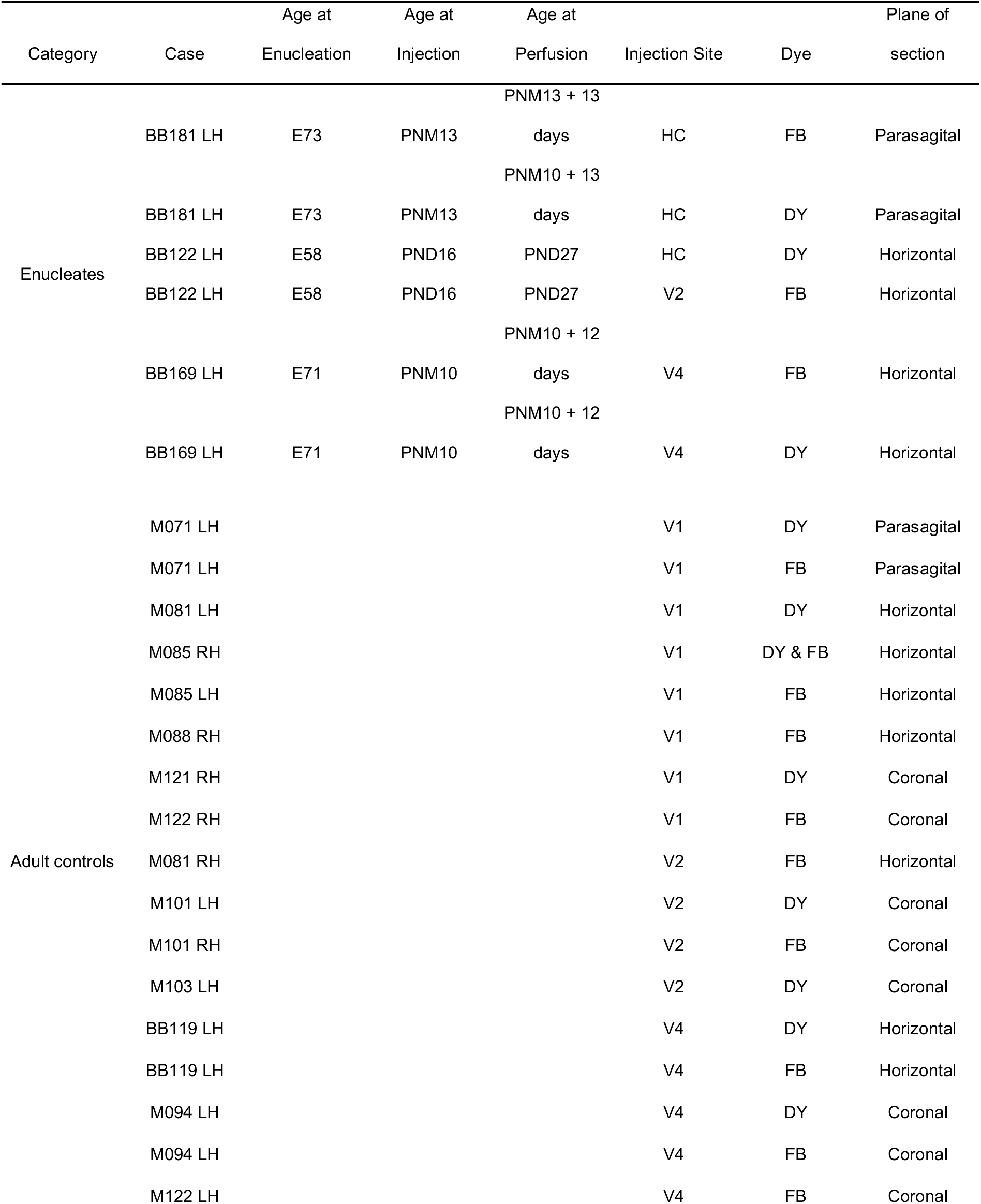
Experimental cases. HC, Hybrid Cortex; E, embryonic day; PND, post-natal day; PNM, post-natal month; DY, Diamidino Yellow; FB, Fast Blue.

| Category | Case | Age at<br>Enucleation | Age at<br>Injection | Age at<br>Perfusion | Injection Site | Dye | Plane of<br>section |
| --- | --- | --- | --- | --- | --- | --- | --- |
| Enucleates |  |  |  | PNM13 + 13 |  |  |  |
|  | BB181 LH | E73 | PNM13 | days | HC | FB | Parasagittal |
|  |  |  |  | PNM10 + 13 |  |  |  |
|  | BB181 LH | E73 | PNM13 | days | HC | DY | Parasagittal |
|  | BB122 LH | E58 | PND16 | PND27 | HC | DY | Horizontal |
|  | BB122 LH | E58 | PND16 | PND27 | V2 | FB | Horizontal |
|  |  |  |  | PNM10 + 12 |  |  |  |
|  | BB169 LH | E71 | PNM10 | days | V4 | FB | Horizontal |
|  |  |  |  | PNM10 + 12 |  |  |  |
|  | BB169 LH | E71 | PNM10 | days | V4 | DY | Horizontal |
| Adult controls | M071 LH |  |  |  | V1 | DY | Parasagittal |
|  | M071 LH |  |  |  | V1 | FB | Parasagittal |
|  | M081 LH |  |  |  | V1 | DY | Horizontal |
|  | M085 RH |  |  |  | V1 | DY & FB | Horizontal |
|  | M085 LH |  |  |  | V1 | FB | Horizontal |
|  | M088 RH |  |  |  | V1 | FB | Horizontal |
|  | M121 RH |  |  |  | V1 | DY | Coronal |
|  | M122 RH |  |  |  | V1 | FB | Coronal |
|  | M081 RH |  |  |  | V2 | FB | Horizontal |
|  | M101 LH |  |  |  | V2 | DY | Coronal |
|  | M101 RH |  |  |  | V2 | FB | Coronal |
|  | M103 LH |  |  |  | V2 | DY | Coronal |
|  | BB119 LH |  |  |  | V4 | DY | Horizontal |
|  | BB119 LH |  |  |  | V4 | FB | Horizontal |
|  | M094 LH |  |  |  | V4 | DY | Coronal |
|  | M094 LH |  |  |  | V4 | FB | Coronal |
|  | M122 LH |  |  |  | V4 | FB | Coronal |

**Table 2.**
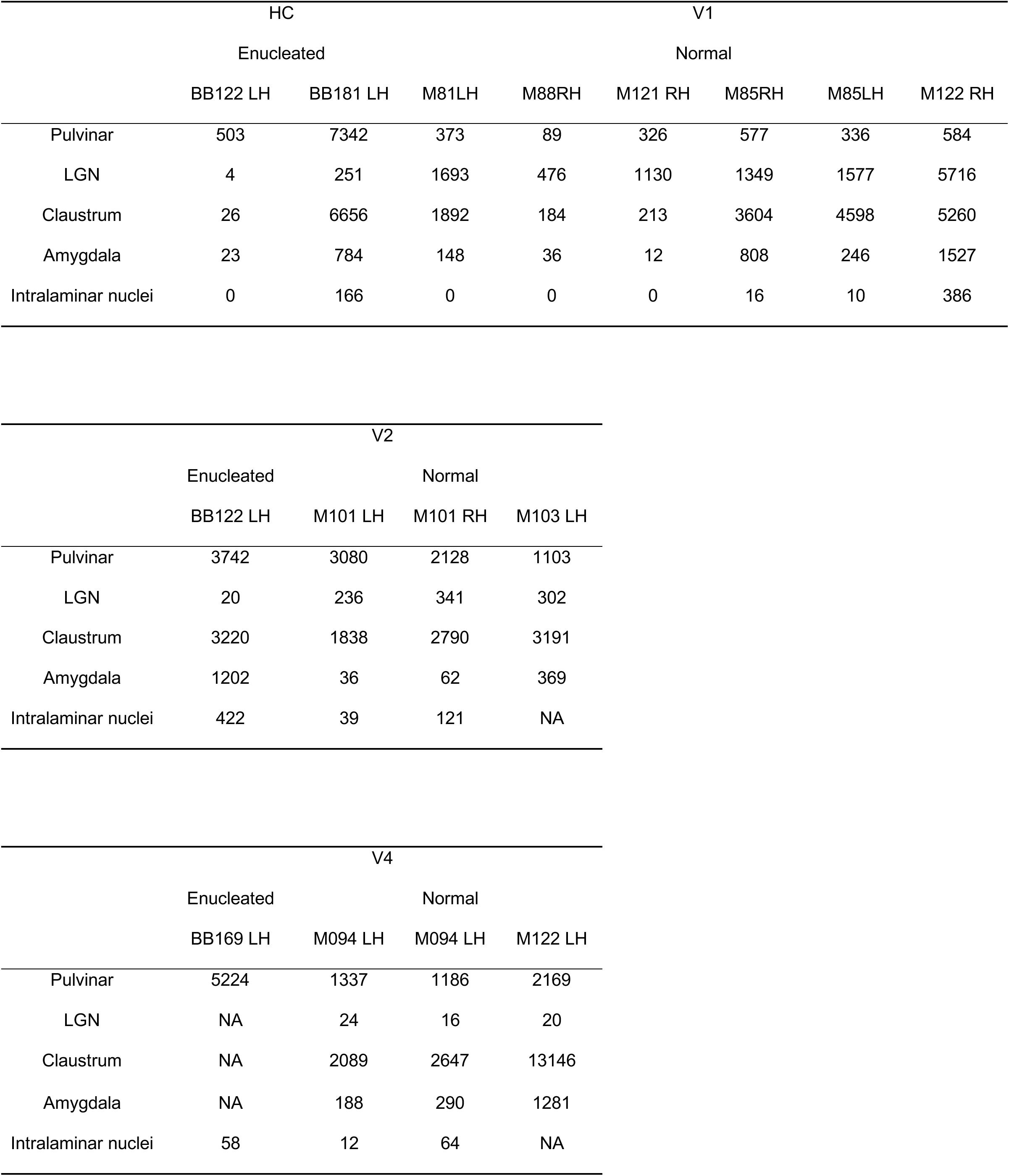
Neuron counts for relevant subcortical structures. Hybrid Cortex, HC; V1, primary visual area; V2, secondary visual area, V4, quaternary visual area.

### Anaesthesia and Surgery

The present study is based on observations following bilateral enucleation performed in three monkey foetuses and contrasted to eight normal controls. The enucleated foetuses were carried to term and after birth injected with retrograde tracers (Diamidino Yellow, DY; and Fast Blue, FB) at different postnatal ages (**Table 1**). Pregnant cynomolgus monkeys (Macaca fascicularis) received atropine (1.25 mg, i.m.), dexamethasone (4 mg, i.m.), isoxsuprine (2.5 mg, i.m.), and chlorpromazine (2 mg/kg, i.m.) surgical premedication. They were prepared for surgery under ketamine hydrochloride (20 mg/kg, i.m) anaesthesia. Following intubation, anaesthesia was continued with 1-2% halothane in a N_2_0/0_2_ mixture (70/30). The heart rate was monitored, and the expired CO_2_ maintained between 4.5% and 6%. Body temperature was maintained using a thermostatically controlled heating blanket. Between embryonic day 58 (E58) and E73 and using sterile procedures a midline abdominal incision was made, and uterotomy was performed. The foetal head was exposed, bilateral eye removal performed, and the foetus replaced in the uterus after closing the incisions. The mother was returned to her cage and given an analgesic (visceralgine, 1.25 mg, i.m.) twice daily for 2 days. All foetuses were allowed normal development until term (E165).

### Injections of Retrograde Tracers

Identical medication, anaesthesia and monitoring procedures were used as described above. Tracer injections were placed in the hybrid cortex, area V2 and area V4. Injections were made by means of Hamilton syringes in a stereotypic fashion. Following injections, artificial dura mater was applied, the bone flaps were closed, cemented and the scalp stitched back into position.

All injections in the anophtalmic brain were confined to the cortical grey matter (Figure 1, S1, S2). Side-by-side injections in target areas of retrograde tracers revealed the topology of connectivity in source areas. Such side-by-side injections were made in the hybrid cortex in the lower part of the medial operculum in case BB181 (**Figure 1**), corresponding in normal cortex to area V1subserving parafoveal visual field (Gattass et al. 1987). In case BB122 a single injection was made in V2 near the lip of the lunate sulcus (**Figure 2**) where foveal visual field is represented in the normal cortex (Gattass et al. 1981). Finally, a pair of very large injections was made on the dorsal part of the prelunate gyrus (**Figure 3**), spanning the central and peripheral representation of area V4 in the normal brain (Li et al. 1989).

**Figure 1.**
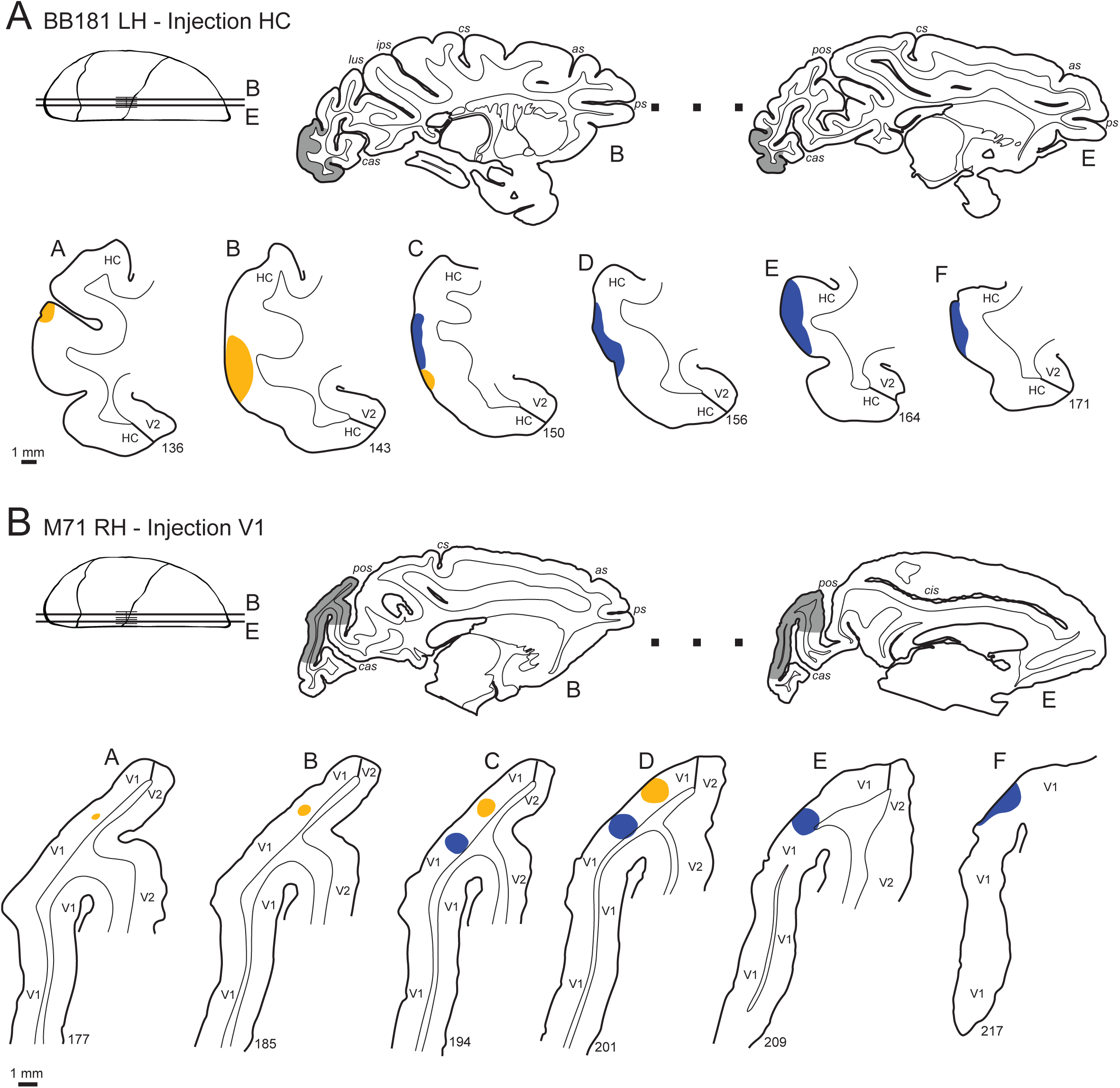
Anatomical drawings showing the injection sites in area V1 in the normal brain and in hybrid cortex in the anophthalmic brain. (A) Tracer injection sites in hybrid cortex in anophtalmic brain (case BB181LH). Upper-row, left cartoons showing location of sections containing injection sites; right low-power view of sections B and E. Bottom-row numbered sections spanning the injection sites. (B) Tracer injection sites in in area V1 in a control (case M071RH). Convention as in (A). Yellow, Diamidino Yellow pick-up zone; blue, Fast Blue pick up zone.

**Figure 2.**
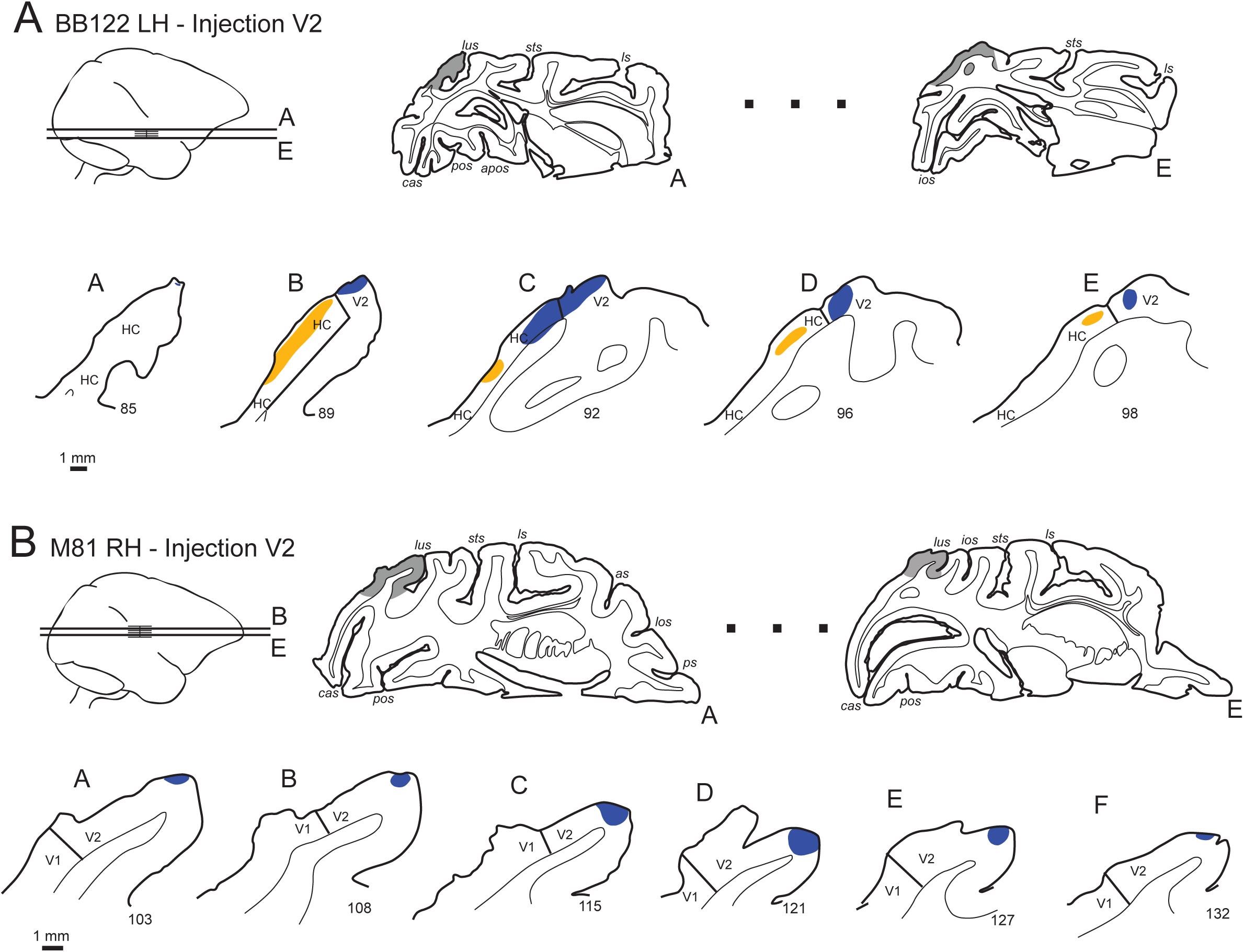
Anatomical drawings showing the injection sites in area V2. (A) Tracer injection sites in area V2 in the anopthalmic brain (case BB122LH). (B) Tracer injection sites in area V2 in a control (case M081RH). Conventions as in Figure 1.

**Figure 3.**
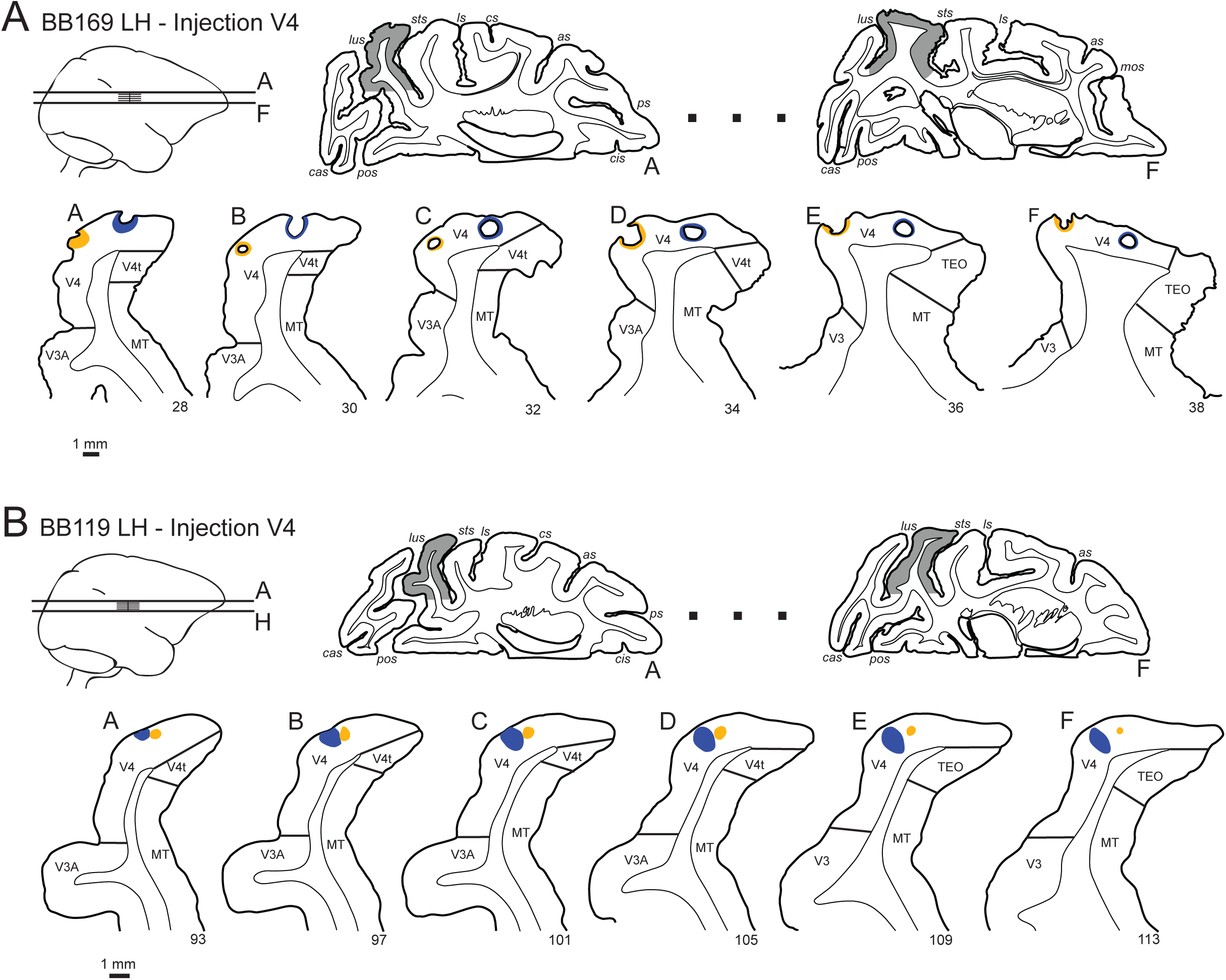
Anatomical drawings showing the injection sites in area V4. **(A)** Tracer injection sites in area V4 in the anophtalmic brain (case BB169LH). (B) Tracer injection sites in area V4 in the normal control (case BB119LH). Conventions as in Figure 1.

The full extent of labelled neurons was charted in subcortical strucutures.

### Animal euthanasia

After 10 to 12 days of recovery that allows optimal retrograde labelling of neurons projecting to the pick-up zone, animals were anesthetised with ketamine (20 mg/kg, i.m.) followed by a lethal dose of Nembutal (60 mg/kg, i.p.) and perfused through the heart with a 1.25% paraformaldehyde and 1.5% glutaraldehyde solution. After fixation, perfusion was continued with a 10-30% sucrose solution to provide cryoprotection of the brain.

### Data Acquisition

Depending on the enucleation case, parasaggital (BB181) or horizontal (BB122 and BB169) sections (40-µm thick) were cut on a freezing microtome and at least 1 in 3 sections were stained for Nissl substance. Controls were cut in horizontal and coronal planes. Sections were observed in UV light with oil-immersion objectives using a Leitz fluorescence microscope equipped with a D-filter set (355-425 nm). High precision maps were made using Mercator software running on Exploranova technology, coupled to the microscope stage. Controlled high frequency sampling gives stable neuron counts despite curvature of the cortex and heterogeneity of neuron distribution in the projection zones of individual areas (Vezoli et al. 2004; Markov et al. 2014b) Characteristics of neurons labeled with FB or DY are described by Keizer and colleagues (Keizer et al. 1983). Area limits and layer 4 were marked on the charts of labeled neurons. These neurons were then attributed to areas using our atlas based on landmarks and histology, and counted according to that parcellation (Markov et al. 2014a).

### Statistical analysis

All statistical analyses were performed in the R statistical environment (R Development Core Team 2016) with additional tools from the MASS, aods3, and betareg packages (Venables and Ripley 2002; Cribari-Neto and Zeileis 2010; Lesnoff and Lancelot 2012). Each injection gave rise to retrogradely labelled neurons, which were plotted and compared against those of normal animals, injected at anatomically equivalent locations. As previously defined (Markov et al. 2011), the FLN (Fraction of Labelled Neurons) is the proportion of cells located in a given source area with respect to the total number of labelled neurons in the cortex. The connectivity profile is defined by the FLN values for each of the structures labelled from the injected target area.

### FLN

The distribution of FLN values has been successfully modelled previously by a negative binomial distribution (Markov et al. 2011; Markov et al. 2014a). This can be performed using a Generalized Linear Model (GLM) with a fixed dispersion parameter (McCullagh and Nelder 1989). In brief, the logarithm of raw neuron counts obtained from each area enter the model as a response variable and the logarithm of the total number of labelled neurons across all areas is used as fixed offset. In this way, the model coefficients estimate FLN values. The dispersion parameter of the negative binomial serves to capture the overdispersion frequently observed in data based on counts (Scannell et al. 2000; Markov et al. 2011). We initially estimated the dispersion parameter for individual areas obtaining values between 2.2 and 7.2, and for subsequent testing used an average value of 4. We then fit this model to compare connection strengths (i.e. FLN values) between normal (i.e. non-enucleated) and enucleated animals. As explanatory variables, we used a 2-level factor, Group (Normal/Anophtalmic) and a 2-level factor, Area for the labelled areas projecting on the target injection. The linear predictor in the GLM includes the main effects of both factors and their interaction. Ninety-five percent confidence intervals were computed to assess the significance of the difference, based on the model fitted estimates.

## Results

In the present study we report on the subcortical labeling observed in the three anopthalmics that showed extensive changes in cortical cytoarchitecture, gross morphology and connectivity that were reported elsewhere (Magrou et al. 2018). Injections of tracer in anophtalmics were aimed at regions housing areas V1, V2 and V4 in 12 normal controls. The increase in the number of controls was in order to increase statistical power in estimations of the quantitative effects of subcortical reorganization in the absence of the retina.

The non-visual thalamus was examined at high magnification in all three anopthalmics for back-labeled neurons. Overall the subcortical structures containing labeled neurons were identical in the experimental and control groups.

### Injection Sites

We describe and quantify the injection sites in controls and experimental cases in order to validate comparing the extent of labeling in subcortical structures. The injections of tracers were made in the region of cortex which is located on the operculum and which in the normal animal corresponds to area V1 but which following early *in utero* removal of the retina assumes hybrid features combining traits observed normally in areas V1 and V2 (**Figure 1**). Injections in presumptive area V2 were made adjacent and posterior to the lunate sulcus (**Figure 2**) and in area V4 on the prelunate gyrus (**Figure 3**).

### Quantitative Changes in Subcortical Distribution of Labeling

In Figures 4, 5 and 6 we compare subcortical labeling in controls and anophtalmics following injections in hybrid cortex in **Figure 4**, area V2 in **Figure 5** and area V4 in **Figure 6**. In the illustration of the results, controls have been chosen such that they have the same plane of section as the experimental animals so as to facilitate comparisons of labeling across cases.

**Figure 4.**
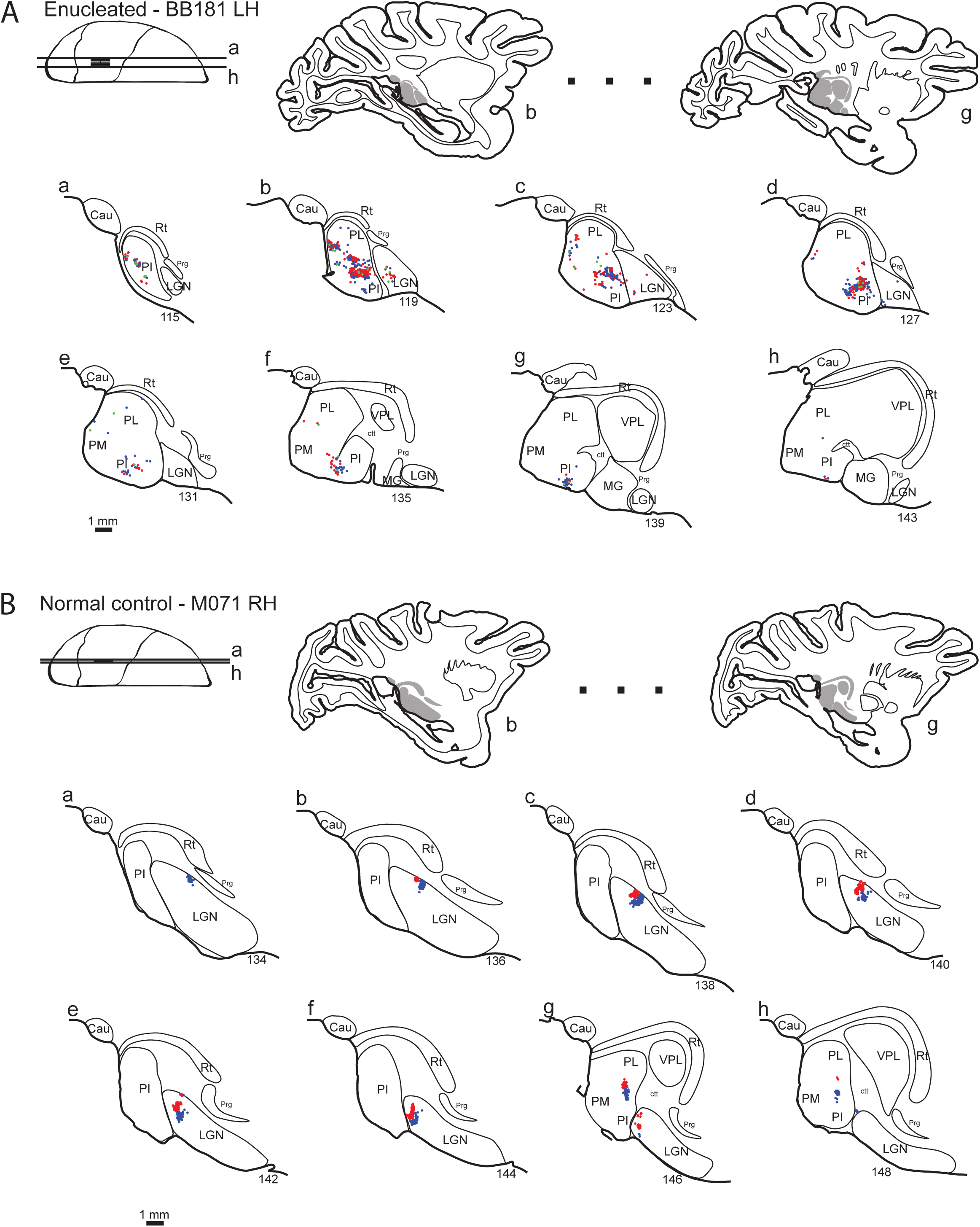
Anatomical drawings showing distributions of labelled cells in subcortical structures following injections in HC/V1. (A) anophtalmic case BB181LH, side-by-side injections in the Hybrid Cortex (HC). **(B)** Corresponding normal control M071RH with a comparable injection site in V1. Blue dots, Fast Blue (FB); red dots, Diamidino Yellow (DY); green dots, doubly labelled cells. For abbreviations of area names see glossary.

**Figure 5.**
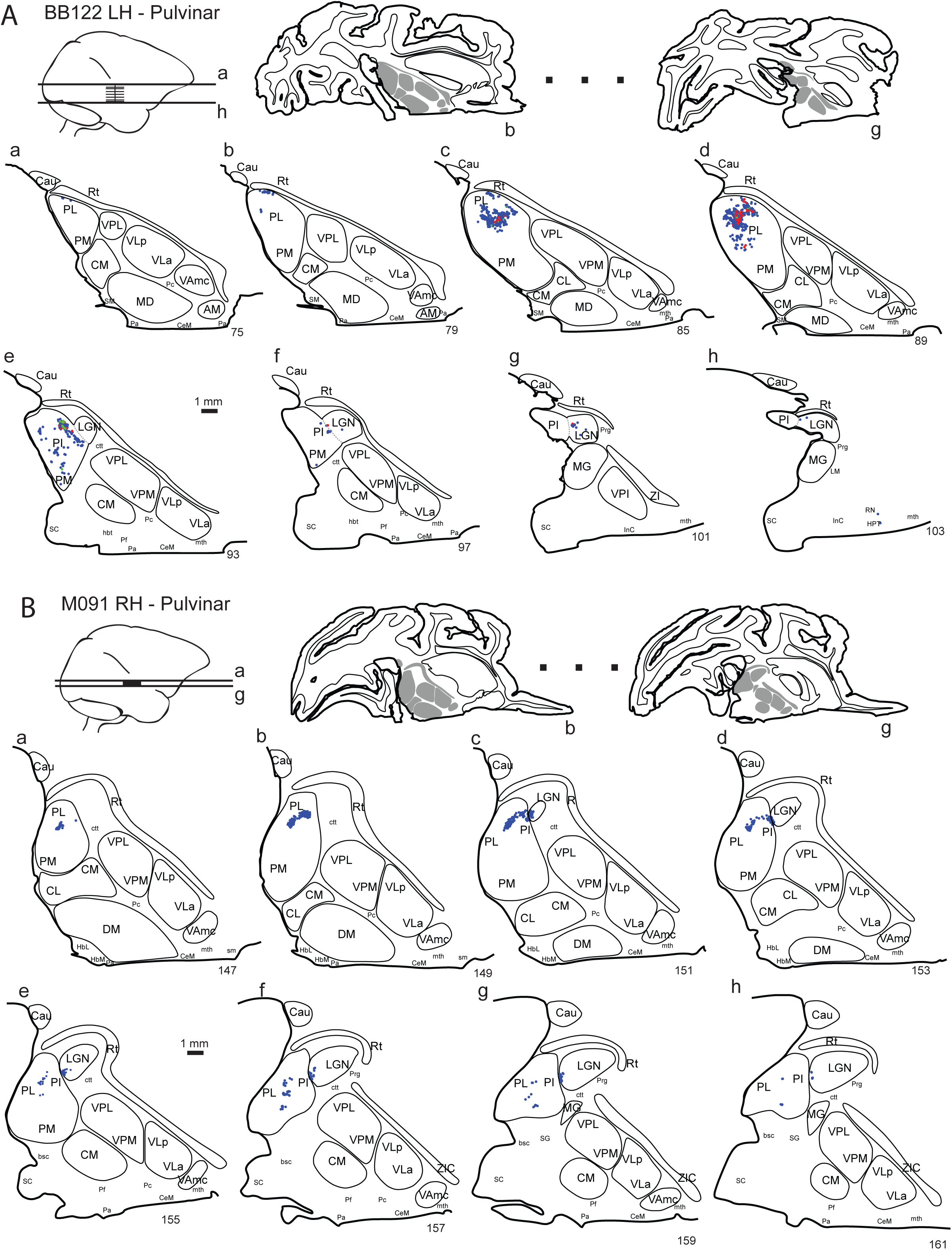
Anatomical drawings showing distributions of labelled cells in subcortical structures following an injection in V2. **(A)** anophtalmic case BB122LH, single injection in V2. **(B)** Corresponding normal control M081RH with comparable injection site in V2. See figure 1 for color coding, for abbreviations of area names see glossary.

**Figure 6.**
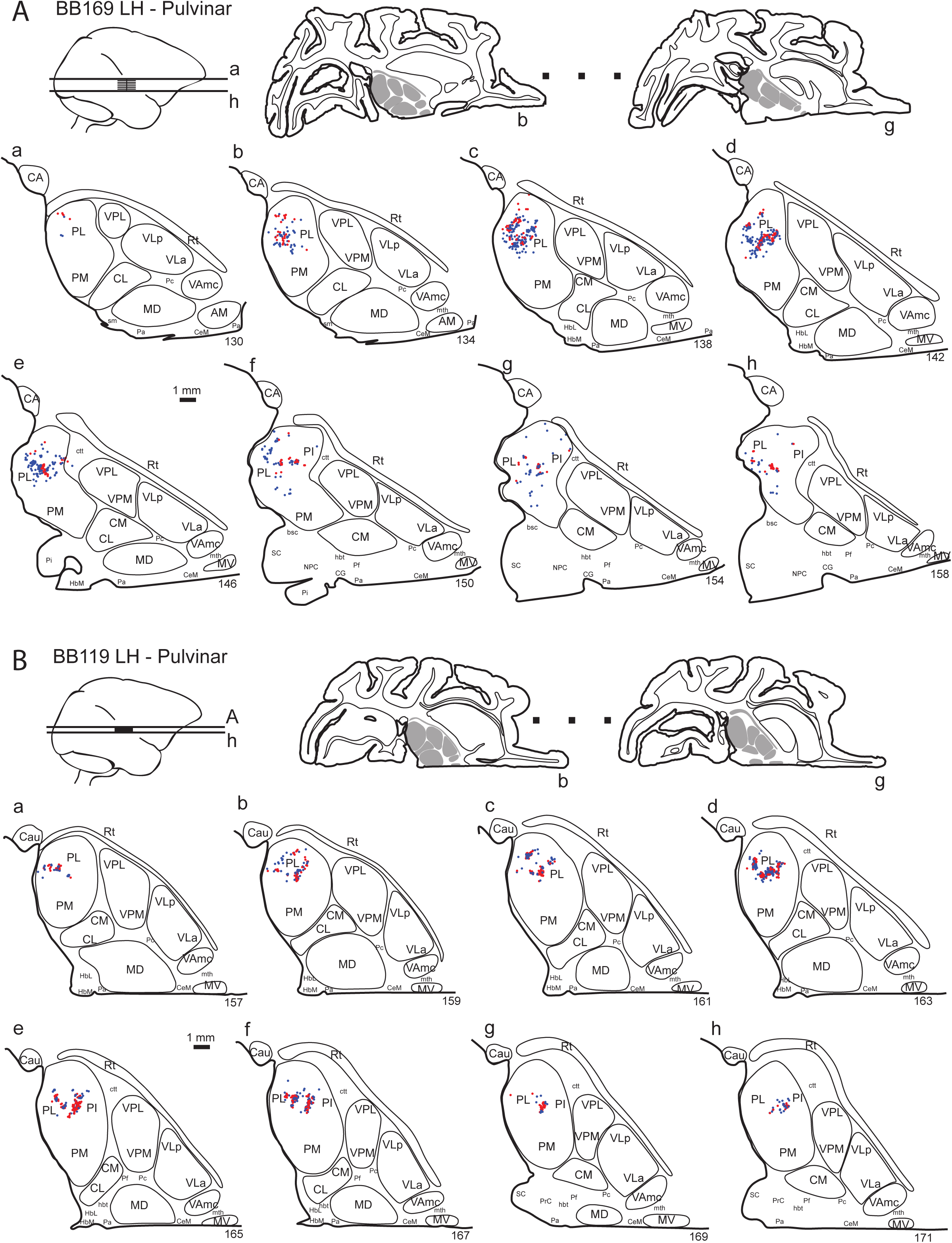
Anatomical drawings showing distributions of labelled cells in subcortical structures following an injection in V2. **(A)** anophtalmic case BB169LH, side-by-side injections in V4. **(B)** Corresponding normal control BB119LH with comparable side-by-side injection sites in V4. See figure 1 for color coding, for abbreviations of area names see glossary.

One anophthalmic animal received side-by-side injections in the hybrid cortex and was sectioned in the parasagittal plane. These injections led to extensive labeling in pulvinar (**Figure 4**), where the Diamidino Yellow and Fast Blue labeled neurons were considerably intermingled, in contrast to the labeling in the normal control where in section 148 in **Figure 4 B**, the two sets of labeled neurons were tightly grouped but segregated. The extent of the labeled neurons in the anophthalmic were very widespread occupying all of the subdivisions of the pulvinar and hence stretching from the most lateral section containing the pulvinar (section 115 **Figure 4A**) to the most medial section (section 143). This strongly contrasted with the control (**Figure 4B**), where the lateral-most regions of the pulvinar from sections 134 to 144 were entirely devoid of labeling where and where labeling was restricted to sections 146 – 148.

Following injection in hybrid cortex, outside of the pulvinar, labeling in the greatly reduced lateral geniculate nucleus was extremely limited in the anophtalmic and comparatively extensive in the normal. In the experimental case, labeled lateral geniculate neurons were concentrated on sections 119, with occasional labeled neuron retching as far as section 127. As in the pulvinar, the two populations of labeled neurons were once again largely intermingled. These observations in the lateral geniculate nucleus of the anophtalmic contrasted with the observation of a much stronger labeling in the lateral geniculate nucleus of the control, with consistent labeling stretching from sections 136 to 146, in which the two populations showed strong segregation forming two adjacent populations on individual sections.

The anophtalmic that received an injection of Fast Blue in area V2 was cut in the horizontal plane and showed strong labeling in the pulvinar largely located in the lateral subdivision of the pulvinar and stretching from sections 75 to section 97 (**Figure 5A**). In this animal there was also a Diamidino Yellow injection in hybrid cortex that largely coincided with the labeling from the area V2 injection, but was more restricted and largely located in sections 85-89. In this animal very few labeled neurons were observed in the lateral geniculate nucleus in sections 93 to 103. Compared to the anophtalmic, labeling in the pulvinar in the control shown in **Figure 5B** was much more compact and large numbers of labeled neurons were found in the lateral-interior subdivisions in the characteristic v shape that has been reported elsewhere (Ungerleider et al. 1983; Kennedy and Bullier 1985).

The anophtalmic that received injections of tracer in area V4 had relatively widespread labeling in the pulvinar (**Figure 6A**). However, the area V4 control shown in **Figure 6B** also showed relatively widespread pulvinar labeling with cells from both injections showing intermingling and it was difficult to define a distinct pattern in the anophtalmic with respect to the control.

### A Selective Increase in Pulvinar Originating Projections

The distributions of labeled neurons reported above suggest that there might be significant changes in the weight of subcortical projections of subcortical inputs to the cortex following the removal of the retina. To address this issue we calculated the FLN for the individual projections (**Figure 7**). This showed a significant increase in the FLN of the pulvinar projection to hybrid cortex compared to the pulvinar projection to area V1 in the normal control. There were no significant differences in the FLN values of the pulvinar projections to either V2 or V4. The projection from the lateral geniculate nucleus was also influenced by removal of the retina: in the enucleate compared to the normal control there was decrease in the projection of the lateral geniculate projection to area V1 and area V2. Are these changes in the visual thalamus echoed by other changes in subcortical structures projecting to the early visual areas? To address this we quantified the FLN in claustrum, amygdala and the intralaminar thalamic nuclei and found no significant differences between control and experimental cases (**Figure 7**).

**Figure 7.**
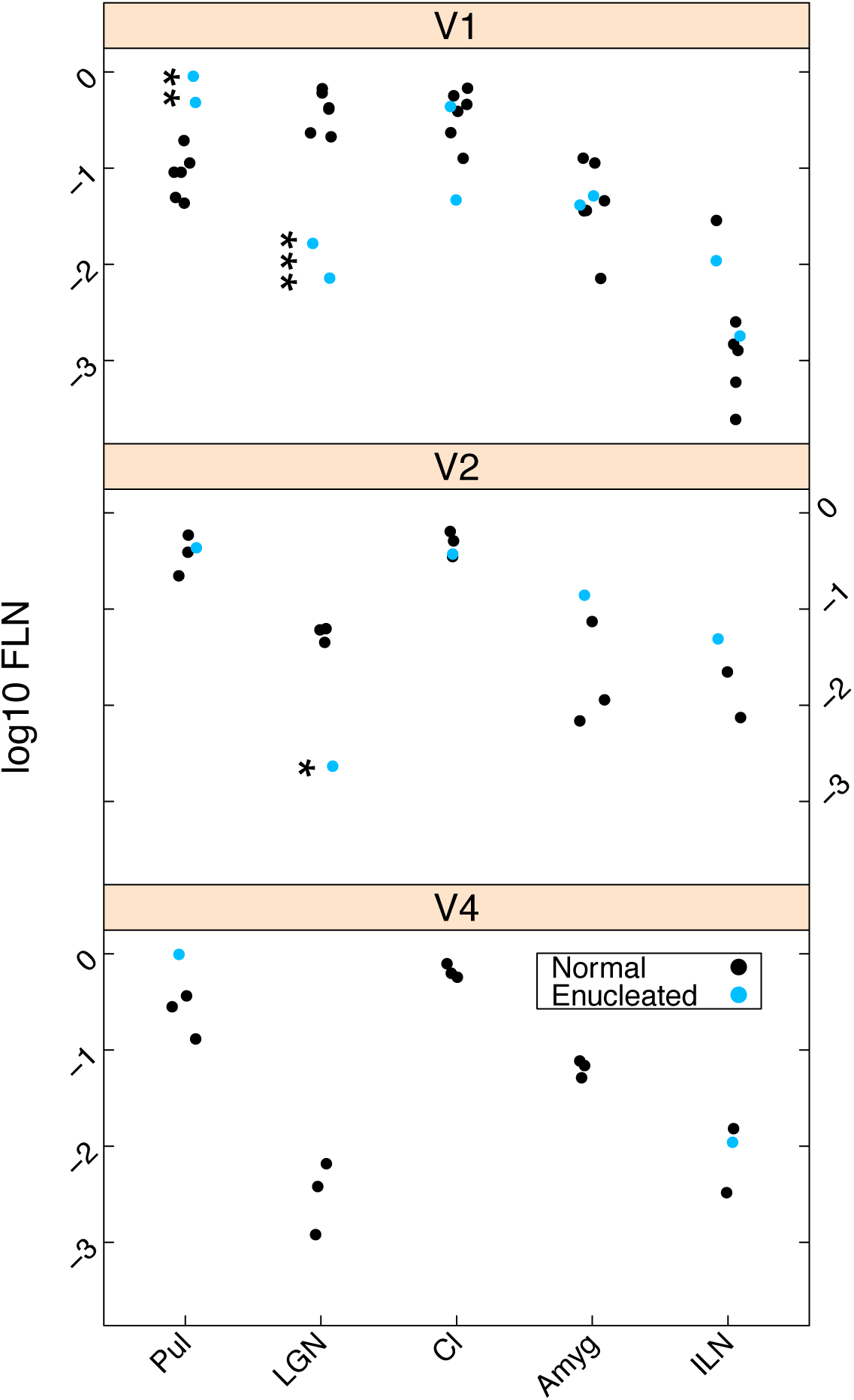
Effect of enucleation on subcortical FLN. Log scale dot plot of FLN. Enucleates, blue dots; normal controls, black dots; upper-panel, injections in normal striate cortex (V1) and Hybrid Cortex (HC) (1anophtalmic, 6 normals); middle-panel, injections in area V2 (1anophtalmic, 3 normals); bottom panel, injections in area V4 (2 enucleates, 3 normal). For abbreviations of area names see glossary.

In order to assess the effect of enucleation on labelling in the pulvinar, illustrated in **Figures 4** and **7**, we estimated the quantitative spread of labelling in that structure. In each section with pulvinar labelling, the smallest polygon encompassing all marked cells was drawn and its area was computed. Volume of labelling for each case was then interpolated from the series of measured. To account for the effect of injection size on the labelling spread, the volume of the uptake zone for each case was also measured and used as a covariate in statistical tests. A linear model (LM) was used to test for differences in pulvinar of the volume of labelling spread between normal controls and anophthalmic animals. The cube root of labelling spread volume was taken as the response variable, following a Box-Cox test to assess the best transformation for normality of the data (Box and Cox 1964; Venables and Ripley 2002). For V1/HC, the difference of spread between normal and anophthalmic groups was significant (p = 0.000126, with a t-value of 6.394 and 9 degrees of freedom (df)), but not for V2 (p = 0.120, t = 2.621, df = 2) and V4 (p = 0.936, t = −0.087, df = 3). These results indicate an effect of anophthalmia on the spread of labelling in the pulvinar for V1/HC but not for V2 and V4. Despite the suggestion in **Figure 5** of a differential spread for V2, the statistical testing failed to attain significance. The enucleated V2 injection, however, is based on a single case and so is likely to be of low statistical power.

Our results show that in the absence of the retina there is a significant increase in the strength of the projection of the pulvinar to the hybrid cortex which is accompanied by a significant increase in the spatial extent of labelling in the pulvinar. The injections in experimental and controls were carried out in a stereotypical fashion. Variations of the uptake zone are not predicted to impact on the FLN values, which are normalised with respect to the total number of labelled neurons. However, variation of the size of the injection and the resulting pickup zone could impact on the spread of labelled neurons that we observe in the pulvinar. In **Figure 8** we compare quantifications of the uptake zones, which showed that up-take zones in the hybrid cortex did not show significant differences.

**Figure 8.**
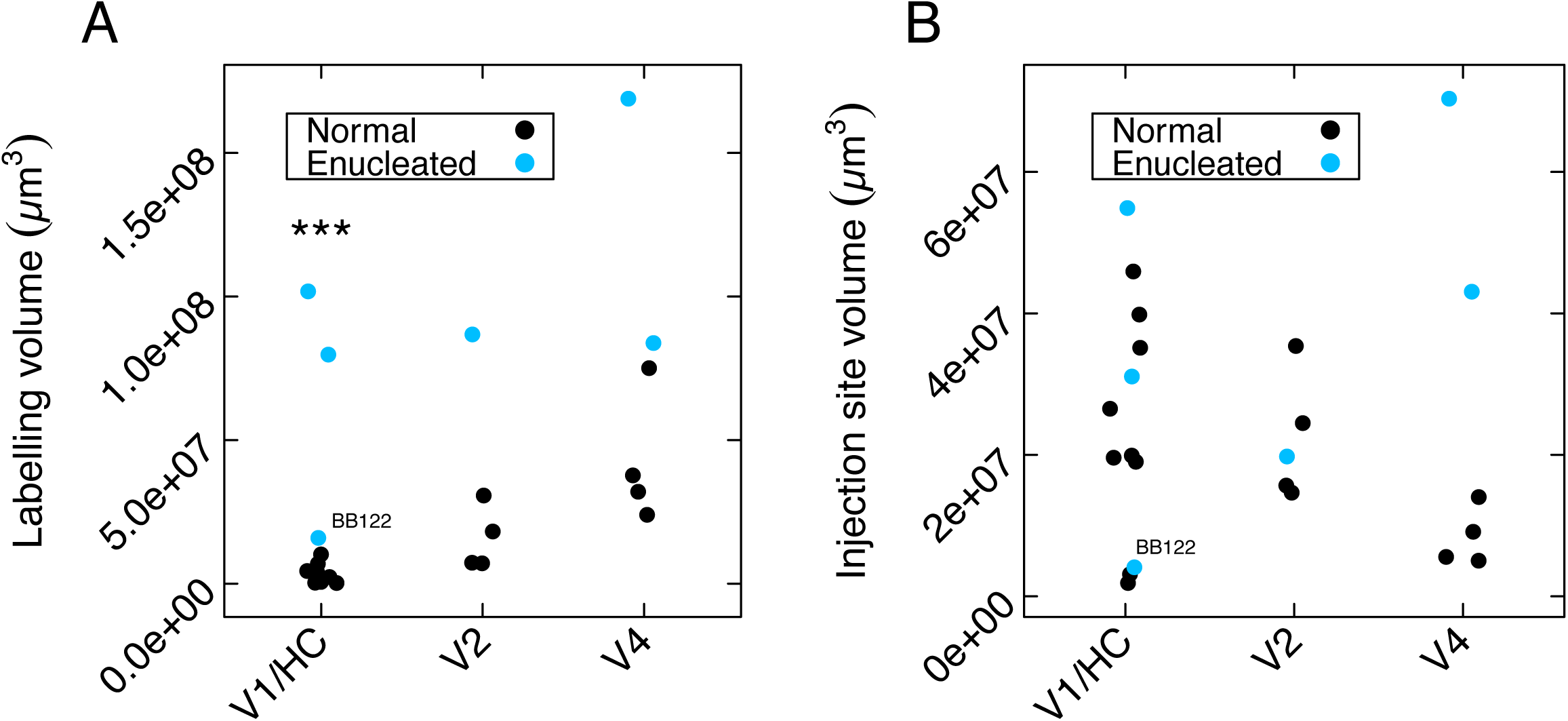
Effect of enucleation on the spread of labelling in the pulvinar. **(A)** Dot plot of the volume of labelling spread in the Pulvinar. **(B)** Dot plot of the volume of the matching injection sites. Enucleates, blue dots; normal controls, black dots. Injections in V1/HC: 3 enucleate, 9 controls; in area V2: 1 anophtalmic, 4 normals; in area V4: 2 anophtalmics, 4 normals. “***” indicates significance with a p-value of 0.000126 (see methods). The case of anophtalmic animal BB122LH is indicated as an outlier for visual reference. For abbreviations of area names, see glossary.

## Discussion

Subcortical labeling in the anophtalmics was quantitatively but not qualitatively different from subcortical labeling in the controls. While the absence of the retina did not lead to the appearance of novel thalamocortical pathways, it did profoundly modify the strength of the projection from the pulvinar complex, which showed significantly increased numbers of labeled neurons with a spatially expanded distribution. This increase in the pulvinar projection to the hybrid cortex was accompanied by a reduction in the projection of the geniculocortical pathway. The expansion of the pulvinar projection to the hybrid cortex appeared to be the unique expansion of ascending pathways as quantification of the projections of the claustrum, amygdala and interlaminar nucleus revealed no differences between experimental and control groups.

### Primate and non-primate models of congenital blindness

There have been a number of investigations in congenitally blind rodents which show many similarities with the present findings. Tracers injected to the visual cortex of congenitally anophthalmic mice found point to point connectivity in the dorsal lateral geniculate nucleus, and while retrogradely labelled neurons were not reported in non-visual thalamus there appeared to be an increase in the incidence of labelled neurons in the lateral posterior thalamic nuclei (Godement et al. 1979; Kaiserman-Abramof et al. 1980). Given that the lateral posterior thalamic nucleus is thought to be evolutionarily related to the pulvinar these results in mutant eyeless mice agree with the present findings. Similar findings have been observed in blind mole rats, animals which have adapted to a non-visual life style (Cooper et al. 1993).

The present findings contrast with those obtained in a recent study on the effects of bilateral enucleation in *Mondelphis domestica*, a South American marsupial (Kahn and Krubitzer 2002; Karlen et al. 2006). Enucleation at post-natal day 4 led followed by tracers injected in area 17 led to retrograde labelling of neurons in subcortical nuclei associated with somatosensory (ventral posterior nuclei), auditory (medial geniculate nuclei), motor (ventrolateral nucleus) systems and limbic/hippocampal systems (anterior dorsal and anterior ventral nuclei(Karlen and Krubitzer 2009). These findings are very different from what is observed in the anopthalmic monkey where subcortical afferents were restricted to the visual thalamus. In a recent study in mouse where geniculocortical pathways were genetically deleted *in utero* likewise failed to find projections to visual cortex from non-visual thalamus (Chou et al. 2013)

The widespread thalamic projections from non-visual thalamus to visual cortex in the anopthalmic marsupial brain (Karlen and Krubitzer 2009), not found in the present study, could be due to the differences in developmental timing with respect to retinal ablation. There are two arguments against this hypothesis. Firstly, while the enucleation at 4th post-natal days in the opossum is at an earlier developmental stage than the enucleation in the macaque at embryonic (E) day 70 (Workman et al. 2013), in both species thalamic innervation of the visual cortex is incomplete and generation of supragranular layers has not yet commenced (Rakic 1974; Robinson and Dreher 1990; Finlay and Darlington 1995; Molnar et al. 1998). Secondly, in both species the ages when the retinae were removed lead to profound and very similar modifications of the cortex, with the generation of a novel cortical region referred to as area X in the marsupial study and hybrid cortex in the present and earlier macaque studies (Rakic 1988; Dehay et al. 1989; Kahn and Krubitzer 2002).

The discrepancy between widespread thalamic projections in the anophtalmic opossum and restricted thalamic projections in the anophtalmic macaque (present study) is difficult to explain. It echoes similar differences found in the cortex where non-visual sensory areas were found to project to visual cortex in the congenitally blind opossum (Karlen et al. 2006) but not in the congenitally blind macaque (Magrou et al. 2018). Elsewhere we have argued that at the level of the cortex these differences could be due to the stabilization of developmental transient projections that have been observed in the opossum (Karlen et al. 2006), and which do not exist in the developing macaque (Magrou et al. 2018). However, such transient widespread projections from different sensory thalamic nuclei have not been reported in anyspecies and therefore cannot be evoked to explain the apparent opossum-macaque differences.

### Developmental specification of the visual pathways

Developmental specification of visual pathways depends on an interplay between genetic and extrinsic mechanisms. Previously, we reported that subsequent to early removal of the retina there were profound modifications of the dimensions, cytology and connectivity of visual cortical areas cortical areas (Magrou et al. 2018). Areal specification is based on morphogens and secreted signaling molecules and extrinsic inputs relayed to the cortex by thalamocortical axons (O’Leary et al. 2007; De la Rossa et al. 2013; Geschwind and Rakic 2013). The role of thalamic axons in arealisation is a multistep hierarchical process involving events at progenitor and neuronal levels (Chou et al. 2013; Pouchelon and Jabaudon 2014; Moreno-Juan et al. 2017). In rodent, genetic depletion of the thalamic input to the primary visual area failed to influence the specificity of thalamic projections to the cortex (Chou et al. 2013). These findings support observations in eyeless mice (Godement et al. 1979) suggesting that matching of thalamic inputs to different sensory areas is relatively genetically constrained. Elsewhere we speculate that differences in primate and rodent on the role of thalamic afferents to the cortex could reflect the early and prolonged thalamic innervation of the germinal zones in the primate compared to the rodent (Magrou et al. 2018).

### Relevance of developmental effects of retinal ablation in macaque on congenital blindness in human

In the anophtalmic macaque the topography of cortico-cortical connectivity and global organization of the ventral and dorsal streams are largely conserved (Magrou et al. 2018), which is also thought to be the case in human congenitally blind (Pietrini et al. 2004; Ptito et al. 2009; Matteau et al. 2010; Striem-Amit et al. 2012a; Striem-Amit et al. 2012b; Striem-Amit et al. 2015; van den Hurk et al. 2017). These findings support an early developmental specification of the functional streams (Deen et al. 2017; Livingstone et al. 2017). Further, in the anophtalmic brain we observe en expansion of the ventral pathway that could reflect cross-modal plasticity (Ptito et al. 2009). However, in addition to the expansion of the ventral stream there are important differences in the organization of the cortex in the congenitally blind macaque model, notably the extensive intrinsic connectivity of hybrid cortex couple to its high location in the cortical hierarchy. The relatively high position in the cortical hierarchy and the conservation of an extensive local connectivity in the hybrid cortex could ensure the long-time constants, which would be required for the observed higher cognitive functions of the deafferentated cortex of the blind (Honey et al. 2012; Chaudhuri et al. 2015; Tomasello et al. 2019).

The present results show that in addition to the extensive reorganization of the cortex there are important changes in the projections to hybrid cortex, namely an important increase in the weight of the projection of the pulvinar and a spatial expansion of cortical projecting neurons in this structure. Note the increase in weight is coherent with findings in eyeless mice, but the greatly expanded spatial distribution is more unexpected (Godement et al. 1979). These findings could have a profound impact on cortical processing given current understanding of the role of information processing by the pulvino-cortical interactions (Arcaro et al. 2018).

The understanding of the dynamic role of the pulvinar in orchestrating communication between cortical areas has come to the fore in recent years (Sherman 2018). There is considerable anatomical evidence that areas that share direct cortical connections are also interconnected via their anatomical connections through the pulvinar (Shipp 2003). These cortical pulvinar loops are thought to play a critical role in active visual processing via gating information outflow (Purushothaman et al. 2012) and synchronizing interconnected areas thereby influencing allocation of attention (Saalmann et al. 2012). The role of the pulvinar in regulating communication between areas is intrinsically tied to the rich interconnectivity with the cortex. The present findings reveal an important expansion of the neurons projecting to hybrid cortex compared to the projection to the normal area V1 (**Figure 4** and 8). Because cortical pulvinar loops directly depend on overlapping of the afferent and efferent connections in the pulvinar, the much-enlarged distribution of pulvinar neurons projecting to the hybrid cortex is expected to allow communication through a much enlarged cortical network in the anophtalmic compared to the controls. Recent work in the Kastner lab suggests that dorsal pulvinar is functionally linked to frontal, parietal and cingulate areas and ventral pulvinar to occipital and temporal regions (Arcaro et al. 2018). Inspection of **Figure 4** shows that whereas in the control there are no labeled neurons in dorsal pulvinar, in the anophtalmic there are numerous labeled neurons in this region. Arcaro et al (Arcaro et al. 2018) showed that the cortical coupling in the dorsal pulvinar is relatively insensitive to low level stimuli and is rather involved in higher order information integration over extended time windows. These findings suggest that the hybrid cortex in the congenitally blind could have an extensive interaction with frontoparietal networks with a broad range of time scales.

This would introduce the possibility for information processing in hybrid cortex being eminently suited to its extensive intrinsic connectivity suggesting it could allow neuronal responses over seconds, which we have speculated would be required for the higher cognitive functions of the deafferented cortex of the blind (Honey et al. 2012; Chaudhuri et al. 2015; Magrou et al. 2018).

## Conclusion

The extensive functional reorganization found in the congenitally blind human cortex has led to the expectation that there will be widespread structural changes with novel, ectopic pathways from different sensory modalities. The macaque model of congenital blindness reported here and elsewhere (Magrou et al. 2018) shows that aside from projections from the subiculum and entorhinal cortex there are no ectopic cortical or subcortical connections to the deafferented cortex. However, given that the interareal network is very dense (Markov et al. 2013), we have taken the precaution of measuring the strength of connections. This revealed major changes both in the cortex and in the subcortical projections. The increase strength and spatial expansion observed in the pulvinar input to the deafferented cortex, along with the modifications that we find in the cortical hierarchy and intrinsic connectivity of hybrid cortex, could have important functional consequences in the blind brain.

## Acknowledgments

This work was supported LABEX CORTEX (ANR-11-LABX-0042) of Université de Lyon (ANR-11-IDEX-0007) operated by the French National Research Agency (ANR) (H.K.), ANR-11-BSV4-501, CORE-NETS (H.K.), ANR-14-CE13-0033, ARCHI-CORE (H.K.), ANR-15-CE32-0016, CORNET (H.K.).

## Author contribution

CD, HK proposed the project; PG, MB, HK surgical intervention; PB, PG, CD histological processing; LM, PB, NTM, GS, HPK, PG, CD data acquisition; LM, PB, NMT, GS, KK, HK data analysis; KK computational modeling; LM, CD, HK wrote the paper.

## Glossary

AM: anteromedial nucleus of the thalamus Amyg, amygdala
Bsc: brachium of superior colliculus
Cau: caudate nucleus
CeM: central medial nucleus of the thalamus
CG: central gray
Cl: Claustrum
CL: central lateral nucleus of the thalamus
CM: central median nucleus of the thalamus
ctt: cortico-tectal tract
HbL: lateral habenular nucleus
HbM: medial habenular nucleus
hbt: habenulopeduncular tract
InC: interstitial nucleus of Cajal
ILN: intralaminar nuclei
LGN: lateral geniculate nucleus
MD: mediodorsal nucleus of the thalamus
MG: medial geniculate complex
mth: mammillothalamic tract
MV: medioventral nucleus of the thalamus
Pa: paraventricular nuclei of the thalamus
Pc: paracentral nucleus of the thalamus
Pf: parafascicular nucleus of the thalamus
PI: inferior nucleus of the pulvinar
Pi: pineal gland
PL: lateral nucleus of the pulvinar
PM: medial nucleus of the pulvinar
PrC: Precomissural nucleus
Prg: pregeniculate nucleus
Pul: Pulvinar
Rt: reticular nucleus of the thalamus
RN: part of the red nucleus
SC: superior colliculus
SG: suprageniculate nucleus
sm: stria medullaris of the thalamus
VAmc: ventral anterior nucleus of thalamus, magnocellular division
VLa: ventral lateral anterior nucleus of thalamus
VLp: ventral lateral posterior nucleus of thalamus
VPL: ventral posterior lateral nucleus of thalamus
VPM: ventral posterior medial nucleus of thalamus
ZIC: zona incerta complex

